# Genetic variation in *ALDH4A1* predicts muscle health over the lifespan and across species

**DOI:** 10.1101/2021.09.08.459547

**Authors:** Osvaldo Villa, Nicole L. Stuhr, Chia-An Yen, Eileen M. Crimmins, Thalida Em Arpawong, Sean P. Curran

## Abstract

Environmental stress can negatively impact organismal aging, however, the long-term impact of endogenously derived reactive oxygen species from normal cellular metabolism remains less clear. Here we define the evolutionarily conserved mitochondrial enzyme ALH-6/ALDH4A1 as a biomarker for age-related changes in muscle health by combining *C. elegans* genetics and a gene-wide association study (GeneWAS) from aged human participants of the US Health and Retirement Study (HRS)^1–4^. In a screen for mutations that activate SKN-1-dependent oxidative stress responses in the muscle of *C. elegans*^5–7^, we identified 96 independent genetic mutants harboring loss-of-function alleles of *alh*-*6*, exclusively. These genetic mutations map across the ALH-6 polypeptide, which lead to age-dependent loss of muscle health. Intriguingly, genetic variants in *ALDH4A1* differentially impact age-related muscle function in humans. Taken together, our work uncovers mitochondrial *alh*-*6/ALDH4A1* as a critical component of normal muscle aging across species and a predictive biomarker for muscle health over the lifespan.

---

Sarcopenia is defined as the age-related degeneration of skeletal muscle mass and is characterized by a progressive decline in strength and performance^8^. This syndrome is prevalent in older adults and has been estimated by large scale studies to afflict 5-13% of people aged 60-70 years and expands to 50% of those aged 80 and above^9^. Loss of muscle function is associated with a decline in quality of life and higher mortality and morbidity rates due to increased chance of falls and fractures^10,11^. Sarcopenia is linked to risk factors, such as a sedentary lifestyle, lack of exercise, and a diet deficient in protein and micronutrients^10^. However, several aspects of the molecular basis of the age-dependent decline in muscle health remains unknown.

In *Caenorhabditis elegans*, mutation of the conserved proline catabolic gene *alh*-*6* leads to premature aging and impaired muscle mitochondrial function^5^. Proline catabolism functions in a two-step reaction, beginning with the conversion of proline to 1-pyrroline-5-carboxylate (P5C) which is catalyzed by proline dehydrogenase, PRDH-1; subsequently, P5C dehydrogenase, ALH-6, catalyzes the conversion of P5C to glutamate. *alh*-*6* mutants have increased levels of P5C; the accumulation of this toxic metabolic intermediate leads to an increase in reactive oxygen species (ROS), including H_2_O_2_^5^, which then activates the cytoprotective transcription factor SKN-1, impairs mitochondrial activity, and drives cellular dysfunction^5–7,12^. When mitochondrial proline catabolism is perturbed, the activation of SKN-1 is restricted to the body wall muscle tissue and only in post-reproductive adults; this is visualized by increased signal in the SKN-1 GFP reporter that targets *gst*-*4*^13^.

While the strong induction of SKN-1 activity in the musculature was linked to mutation of the mitochondrial P5C dehydrogenase gene, and not observed in other genetic mutants^6,7,12,14–17^, the breadth of genetic mutations that could induce SKN-1 activation in muscle was unknown. In order to identify additional genetic components of this age-related muscle phenotype, we performed an ethyl methanesulfonate (EMS) mutagenesis screen selecting for the same age-dependent activation of the *gst*-*4p::gfp* reporter in the musculature. We screened the progeny of ~4000 mutagenized F1 animals and isolated 96 mutant animals with age-dependent activation of the *gst*-*4p::gfp* reporter restricted to the body wall musculature, which phenocopies the *alh*-*6(lax105)* mutant (**Fig. 1a**, Supplementary Fig. 1). To rule out additional loss-of-function alleles of *alh*-*6* we performed genetic complementation (cis-trans) testing with our established *alh*-*6(lax105)* allele; surprisingly, all 96 new mutations failed to complement and as such were all loss-of-function alleles of *alh*-*6*.

**Figure 1.**
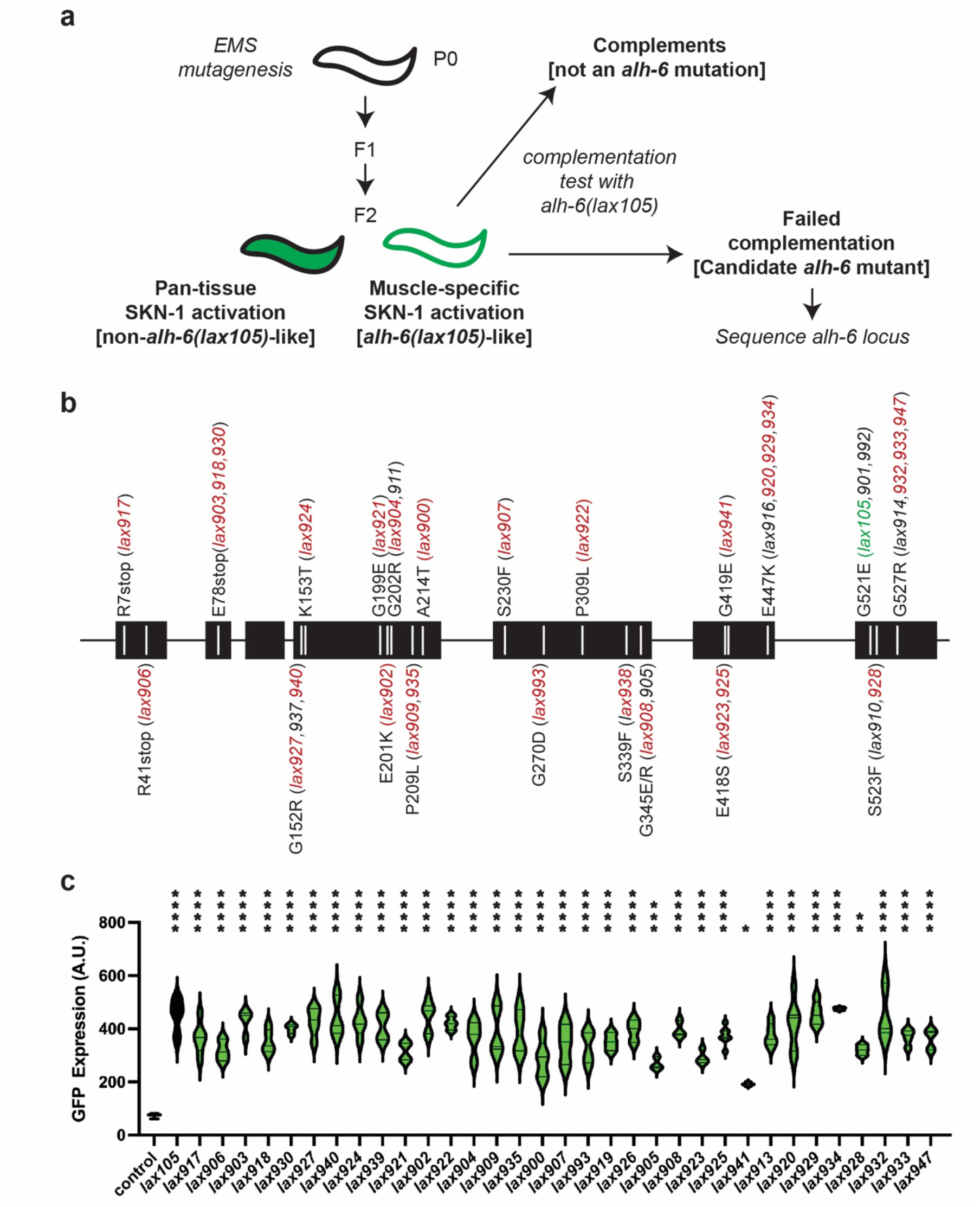
Mutation of *alh*-*6* uniquely activates age-dependent and muscle-specific activation of SKN-1 cytoprotection. (**a**) Schematic representation of genetic screen for mutants that phenocopy *alh-6(lax105)*. (**b**) Molecular identity of mutants isolated and sequenced. Alleles that were selected for additional functional tests of muscle function (**Fig. 4**) are highlighted in red and the location of the canonical *alh*-*6(lax105)* allele is highlighted in green. (**c**) Quantification of SKN-1 activation in the muscle in the new *alh*-*6* mutant alleles, as measured by the intensity of GFP fluorescence from the SKN-1 reporter *gst*-*4p::gfp* (see Supplemental Figure S1 for representative images). One-way ANOVA relative to control (wildtype *alh*-*6*) animals; *, p<0.05; **, p<0.01; ***, p<0.001; ****, p<0.0001.

To catalog these mutations, we began sequencing the *alh*-*6* genomic locus in each of the mutants isolated. After sequencing approximately half of the mutants, we noted the repeated independent isolation of several distinct molecular lesions in *alh*-*6*: E78Stop (*lax903*, *lax930*), E447K (*lax916*, *lax920*, *lax929*, *lax934*), G527R (*lax914*, *lax932*, *lax933*, *lax947*), etc. (**Fig. 1b**). The lack of diversity in genes uncovered and the independent isolation of identical alleles multiple times from this unbiased screen suggests genetic saturation and specificity of this response to animals with defective mitochondrial proline catabolism. In addition, several mutations mapped to discreet regions of the linear ALH-6 polypeptide, including G152/K153, G199/E201/G202, and E418/G419, which may define critical domains in the folded protein. Imaging at day 3 of adulthood revealed that each mutant was phenotypically identical to *lax105* in the localization of SKN-1 activation (Supplementary Fig. 1), but with varying intensity (**Fig. 1c**). Taken together, these data reveal that the age-dependent and muscle-restricted activation of SKN-1 is driven specifically by mutations in mitochondrial *alh*-*6*.

Based on the striking specificity of the muscle-restricted and age-dependent activation of SKN-1 in *C. elegans* harboring mutations in *alh*-*6*, combined with the high degree of conservation in mitochondrial metabolism pathways across metazoans^6^, we reasoned that *ALDH4A1* genetic variants would associate with muscle phenotypes of human aging. To test this hypothesis, we performed gene-wide association scans (GeneWAS) adjusting for relevant covariates and indicators of population stratification in the US Health and Retirement Study (HRS); a nationally representative longitudinal study of >36,000 adults over age 50 in the US^2,18^. HRS collects biological and genetic samples on sub-sets of participants and assesses physical and psychosocial measures of all study participants over adulthood; including multiple measures of muscle function (Supplementary Table 1). We identified 53 single nucleotide polymorphisms (SNPs) within the *ALDH4A1* region that were associated with age-related muscle function, out of the 273 possible variants on the HRS array (**Table 1** and **Fig. 2**). Overall, four associations between variants within *ALDH4A1* and phenotypes were detected and surpassed the gene-wide significant cut-off of *p*≤0.003, corrected for multiple-testing; they were observed between three specific SNPs in *ALDH4A1* (**Table 1**). Additionally, 13 associations were observed between variants within *ALDH4A1* and phenotypes that exceeded the *p* ≤ 0.013 cutoff for a suggestive association; this occurred between nine specific SNPs and eight phenotypes (**Table 1**).

**Table 1.** Top SNPs associated with specific phenotypes

| Phenotype | SNP name | Location | Ref. Allele | Minor allele | Freq.<sup>1</sup> | Scan N<sup>2</sup> | Effect Size<sup>3</sup> | P-value |
| --- | --- | --- | --- | --- | --- | --- | --- | --- |
| ***Gene-wide Significance Level*** |  |  |  |  |  |  |  |  |
| Walking across a room | rs111289603 | 19195492 | G | G | 0.017 | 9907 | 0.399 | 4.5E-04 |
| Grip strength decline | rs28665699 | 19200185 | A | A | 0.014 | 5228 | -0.045 | 9.1E-04 |
| IADL decline | rs111289603 | 19195492 | G | G | 0.017 | 9041 | 0.032 | 2.4E-02 |
| Gait speed decline | rs77608580 | 19196968 | A | G | 0.017 | 3319 | 0.052 | 2.5E-03 |
| ***Suggestive Level*** |  |  |  |  |  |  |  |  |
| Mobility decline | rs28665699 | 19200185 | A | A | 0.014 | 8978 | -0.028 | 4.1E-03 |
| Grip strength ability | rs2230707 | 19202896 | A | G | 0.484 | 8845 | 0.018 | 7.6E-03 |
| Gait speed decline | rs28756052 | 19203375 | A | A | 0.280 | 3327 | 0.046 | 7.6E-03 |
| Gait speed decline | rs11740 | 19199221 | A | T | 0.471 | 3331 | -0.046 | 8.4E-03 |
| Lifting 10 pounds | rs28493067 | 19203333 | A | T | 0.473 | 9583 | 0.092 | 9.8E-03 |
| Walking 1 block | rs111289603 | 19195492 | G | G | 0.017 | 9842 | 0.239 | 1.1E-02 |
| Walking across a room | rs113232075 | 19211163 | G | G | 0.014 | 9900 | 0.353 | 1.2E-02 |
| Gait speed decline | rs61749348 | 19200988 | A | A | 0.065 | 3332 | 0.043 | 1.3E-02 |
| Gait speed decline | rs28594104 | 19202282 | A | T | 0.078 | 3330 | 0.042 | 1.5E-02 |
| Getting up from a chair | rs111289603 | 19195492 | G | G | 0.017 | 9882 | 0.200 | 1.5E-02 |
| Mobility decline | rs28493067 | 19203333 | A | T | 0.473 | 8980 | 0.024 | 1.5E-02 |
| Lifting 10 pounds | rs16862341 | 19220748 | C | G | 0.427 | 9625 | -0.082 | 1.6E-02 |
| IADL2 decline | rs111289603 | 19195492 | G | G | 0.017 | 9041 | 0.023 | 1.7E-02 |

1. Freq = frequency of minor allele as reported by 1000 genomes.
2. N = Sample size of scan for the phenotype and SNP.
3. Effect sizes provided are standardized regression coefficients.

**Figure 2.**
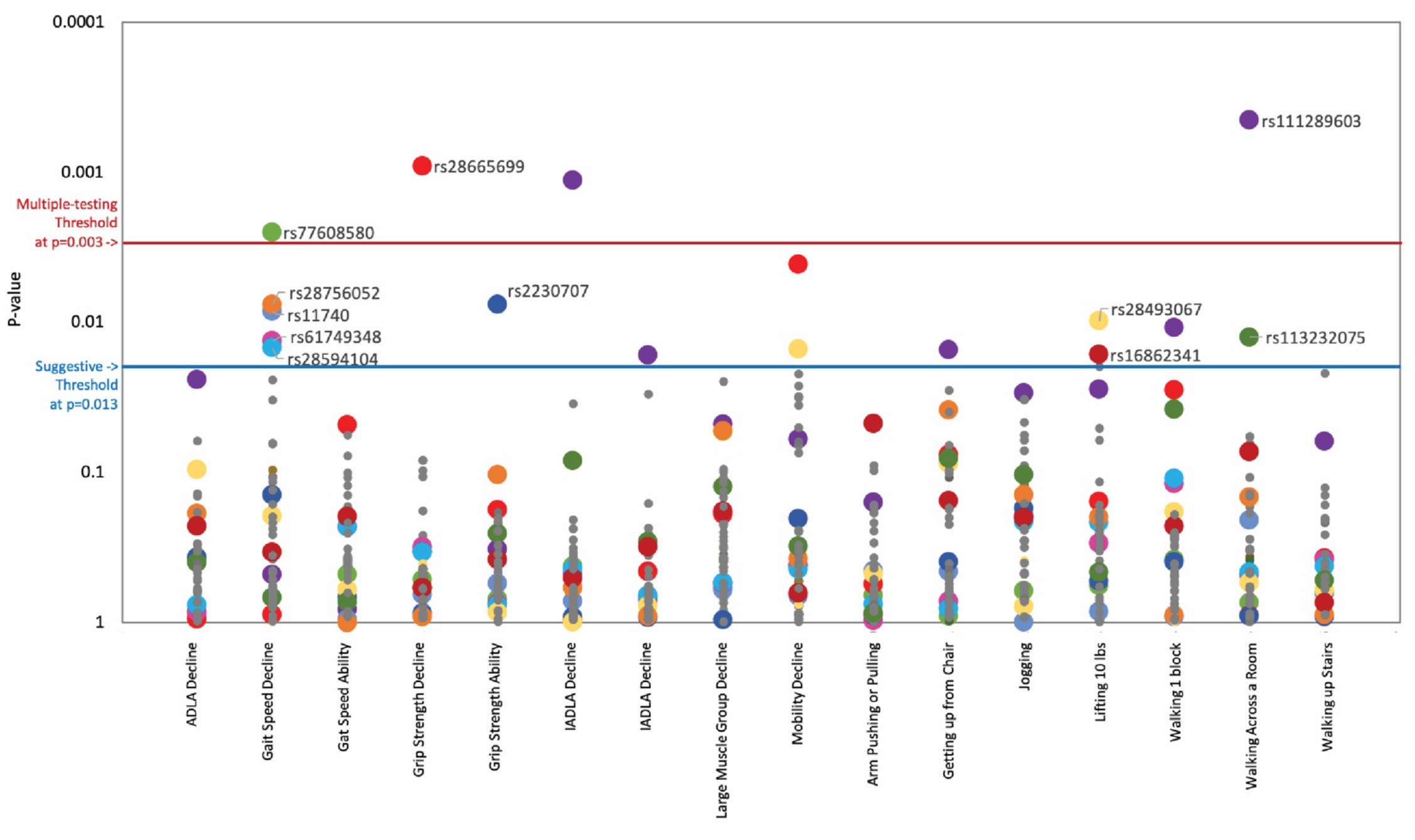
*ALDH4A1* variants associate with multiple human age-related phenotypes of muscle function. Plot of association between variants in the *ALDH4A1* gene and aging-related muscle function and decline in the U.S. Health and Retirement Study (HRS). The x-axis shows each of the phenotypes arranged in order of primary interest. The y-axis shows the log of the p-value for the association between the SNP and the phenotype. The top SNPs across phenotypes are coded by color and size, with larger dots identifying SNPs that surpassed the gene-wide or suggestive thresholds for at least one phenotype. Only the top association for each SNP is labeled with the SNP name. To identify where these same SNPs fall in association with different phenotypes, the top SNPs are coded with the same color across phenotypes.

We calculated phenotypes for aging-related muscle decline over time because they are more robust for testing genetic associations and representing aging processes than single-time point assessments of muscle functioning. Three SNPs associated independently with more than one aging-related muscle phenotype (Supplementary Table 2). Difficulty with performing activities of daily living (ADL) and instrumental activities of living (IADL) are strongly associated in older populations with poor muscle measures^19^. Five of the muscle aging phenotypes reflect individual abilities, whereas three reflect composite scores of functioning (e.g., IADL decline, mobility) (Supplementary Table 1). These results indicate that variants within the *Aldh4a1* locus affect an individual’s ability to perform basic ambulatory movements such as walking short distances and getting up from a sitting position.

We identified patterns of overlap for SNPs associated with more than one phenotype with varying strength (**Fig. 2**). For rs111289603, we calculated the association with the phenotype of having difficulty walking across a room as having an odds ratio (OR) = 1.49, 95% Confidence Interval of 1.19 to 1.86 (beta = 0.40, SE = 0.11, *p*-value = 0.00045). This is interpreted as an additive effect such that for each additional minor allele (G allele) for SNP rs111289603, on average, individuals have a 49% increase in odds for having difficulty walking across a room compared to their same aged counterparts who do not have the effect allele. rs77608580 was significantly associated with change in gait speed over time (**Fig. 3a**). Specifically, with each additional A allele, there was an average increase in gait speed of 0.052 meters per second per year compared to other same-aged individuals without the allele (*p*-value = 0.0025). This was assessed among N=3,319 older individuals with a mean age of 73.0 years (SD=5.9) and mean gait speed of 0.80 meters per second (SD=0.25), or 2.6 ft/sec. Five SNPs located within 5,314 bps of each other (rs77608580, rs28756052, rs11740, rs61749348, and rs28594104) are associated with decline in gait speed in this analysis, one at the gene-wide significance level and four at the suggestive level (**Fig. 2**).

**Figure 3.**
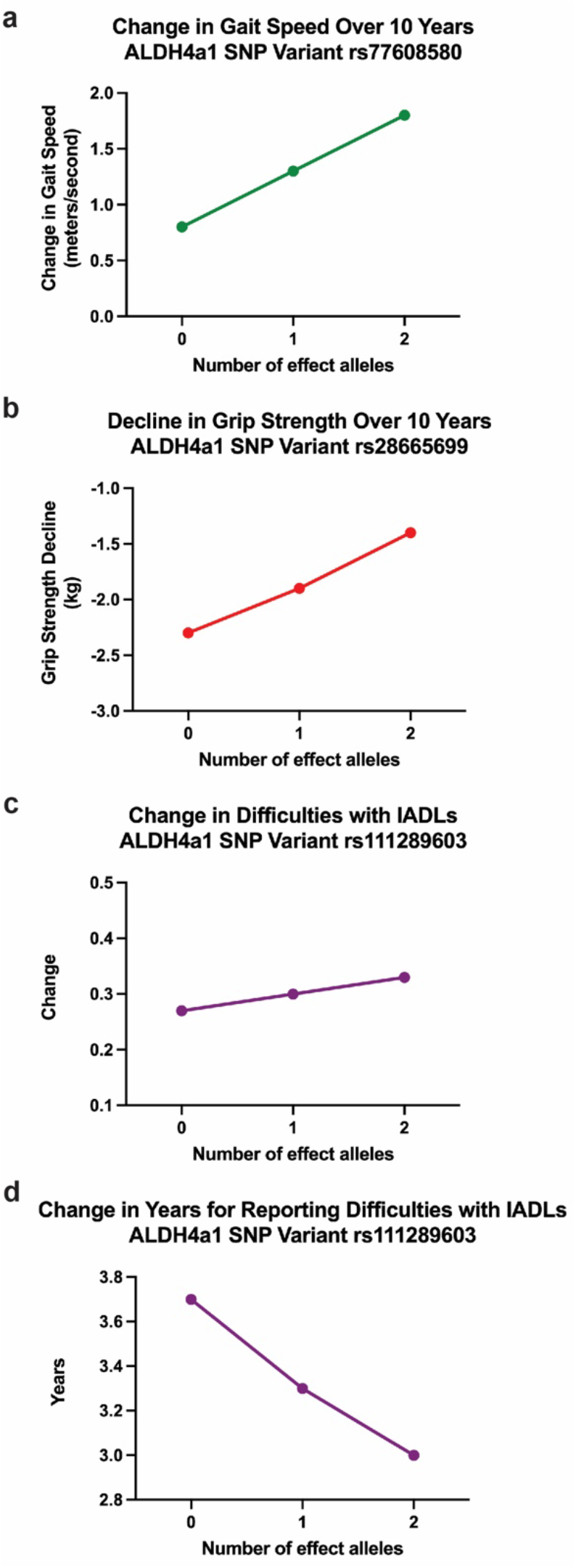
Effects of *ALDH4A1* variation on muscle function with age. (**a**) Change in gait speed over 10 years. Effect of SNP rs77608580 on aging-related changes in gait speed (*b*=0.052, p=0.0025). Over the span of 1 decade, on average, those with 1 or 2 effect alleles will have faster gait speeds with a difference of 0.52 and 1.04 m/sec, respectively, compared to those without an effect allele. (**b**) Decline in grip strength over 10 years. Variation in *ALDH4A1* (SNP rs28665699) is inversely associated with decline in aging-related grip strength (*b*=-0.045, p=0.0009). Individuals with 1 or 2 effect alleles have slower progression of weakened grip strength over 10 years by 0.5 and 1.0 kg respectively, compared to the same aged individuals without the effect allele. (**c**) Change in difficulties with IADL. Effect of variation in *ALDH4A1* (SNP rs111289603) on the rate for reporting difficulties in performing instrumental activities of daily living (IADL) with increasing age (*b*=0.032, p=0.001). (**d**) Change in years for reporting difficulties with IADL. Difference in the number of years for reporting difficulties with performing an IADL associated with each effect allele in rs111289603.

Measures of muscle health, such as grip strength, are effective biomarkers of overall health in older populations^20,21^. rs28665699 was significantly associated with an increase in grip strength over time; with each additional A allele, there was an average increase in grip strength by 0.045 kg weight per year while holding all other characteristics constant (age, sex, and use of the dominant hand for gripping; *p*-value = 0.0009). This was assessed among N=5,228 older individuals with a mean age of 68.9 years (SD=10.4), mean grip strength of 30.21 kg (SD=11.1), and average level of decline in grip strength at 2.31 kg per year (SD=5.37). If calculated as change over a ten-year period, those with 1 or 2 effect alleles would have stronger grip by 0.5 and 1.0 kg compared to those without an effect allele, respectively. The allele therefore is associated with a slower rate of decline in grip strength over a decade of age (**Fig. 3b**).

Instrumental activities of daily living (IADL) are defined by the ability to perform tasks of using a telephone, taking medication, and handling money, all of which are sensitive to muscle function. rs111289603 was significantly associated with difficulties with performing IADL over time such that with each additional G allele, there was a predicted increase in difficulty by 0.032 units per year (*p*-value = 0.001), while holding all other characteristics (age, sex) constant (**Fig. 3c**). For older individuals without the effect allele, it would take an average of 3.7 years before reporting difficulty with performing an additional IADL task (i.e., from 0 difficulty to 1, or 1 to 2, or 2 to 3 difficulties) whereas for those with 1 or 2 effect alleles, it would take an average of 3.3 or 3.0 years, respectively, before reporting difficulty with an additional task (**Fig. 3d**, N=9041 older individuals).

It is not known if any one identified *ALDH4A1* SNPs is a causal variant or if they mark a different variant within the *ALDH4A1* gene that was not represented on the HRS array. Regardless, the consistency of these results collectively supports a true association between *ALDH4A1* and age-related muscle function. To test how genetic variation in P5C dehydrogenase can influence age-related muscle function, we returned to our collection of *C. elegans* strains harboring mutations in *alh*-*6*. We measured individual animal movement speed as a function of muscle health with age^22^. Only animals harboring mutations of ALH-6 at position G152R(*lax940*), K153T(*lax924*), S523F(*lax928*), and G527R(*lax933*) resulted in a significant loss of movement speed at larval stage 4; just prior to adulthood (**Fig. 4a**). However, with the exception of S230F(*lax907*) and Y427N(*lax918*), all mutants tested displayed a significant reduction in movement speed at Day 3 of adulthood (**Fig. 4b**). Taken together these data support the age-specific acceleration of muscle decline in mitochondrial proline catabolism mutants, which is conserved from nematodes to humans.

**Figure 4.**
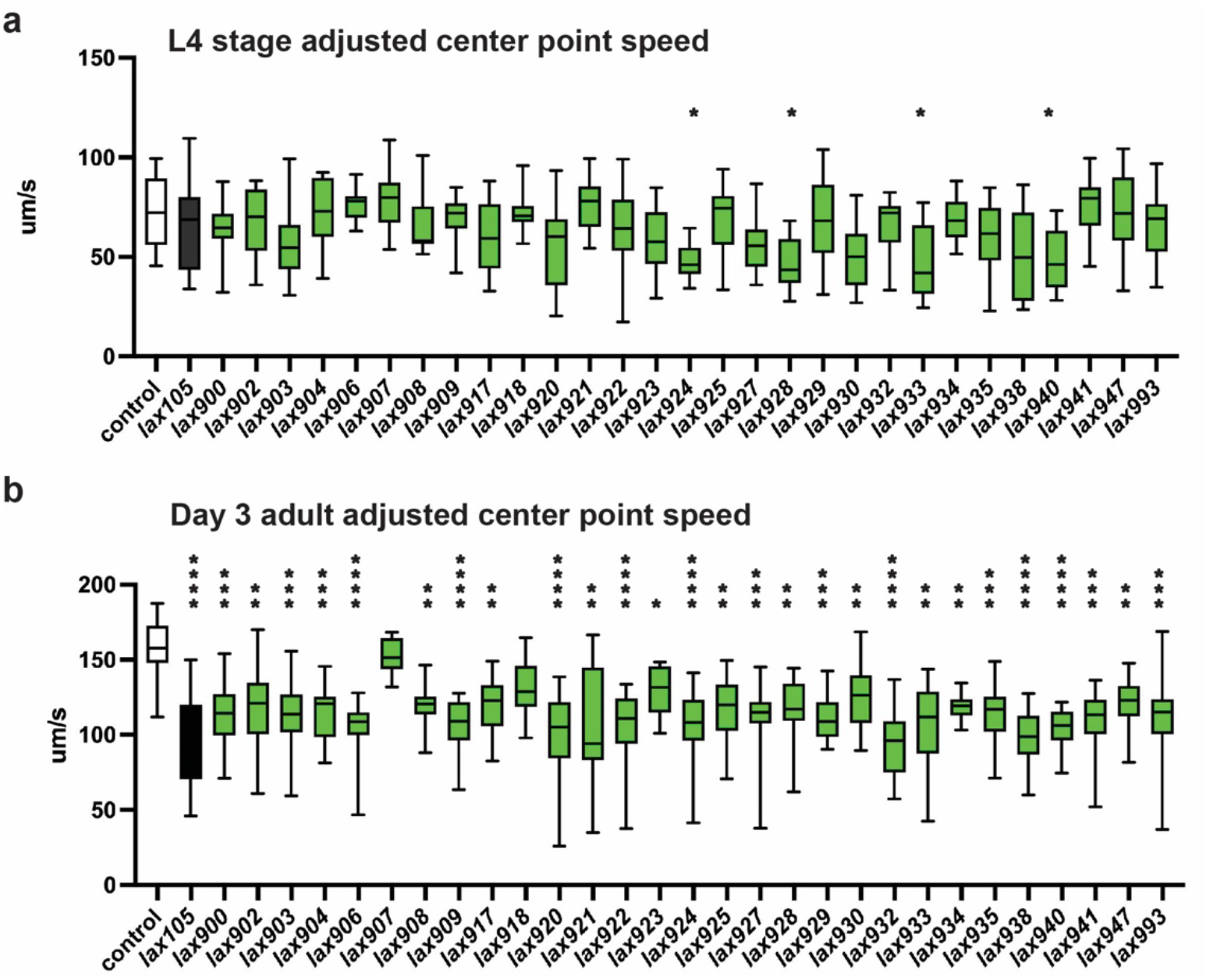
*alh*-*6* mutations accelerate loss of muscle function. WormLab software analysis of adjusted center point speed of individual animals of the given genotypes at the L4 stage (**a**) or day 3 of adulthood (**b**). Brown-Forsythe and Welch ANOVA test with Dunnett’s T3 multiple comparisons test, with individual variances computed for each comparison. *, p<0.05; **, p<0.01; ***, p<0.001; ****, p<0.0001.

Existing biomarkers of muscle health that can accurately predict muscle health later in life are extremely scarce due to limited data in human aging and an incomplete understanding of the molecular basis of sarcopenia. Understanding the diversity of genetic variation underlying sarcopenia, as well as their corresponding phenotypic outcomes, will be critical for providing accurate risk assessments for family planning and genetic counseling of older adults. Although the etiology of human disease is complex and multifactorial, we have used a combination of classical *C. elegans* genetics and human genetic association studies to define genetic variation in *alh*-*6/ALDH4A1* as a new biomarker of age-related muscle health in human.

## ACKNOWLEDGEMENTS

We thank J. Gonzalez for technical assistance and Drs. R. Irwin and C. Duangjan for critical reading of the manuscript. Some strains were provided by the CGC, which is funded by the NIH Office of Research Infrastructure Programs (P40 OD010440). This work was funded by the NIH AG058610 and AG063947 to S.P.C., T32AG052374 to O.V. and N.L.S. and T32GM118289 to N.L.S. This study was supported in part by funding from The National Institute on Aging, through the USC-Buck Nathan Shock Center (P30 AG068345). The National Institute on Aging has supported the collection of both survey and genotype data for the Health and Retirement Study through co-operative agreement U01 AG009740. The datasets are produced by the University of Michigan, Ann Arbor. The HRS phenotypic data files are public use datasets, available through: https://hrs.isr.umich.edu/data-products/ access-to-public-data. The HRS genotype data is available to authorized researchers: https://www.ncbi.nlm.nih.gov/projects/gap/cgi-bin/study.cgi?study_id=phs000428.v2.p2

## AUTHOR CONTRIBUTIONS

S.P.C. designed the study; O.V., C-A.Y., N.L.S., T.E.A., and S.P.C. performed the experiments; O.V., C-A.Y., N.L.S., E.M.C., T.E.A., and S.P.C. analyzed data. S.P.C. wrote the original manuscript and all authors revised the final manuscript.

## METHODS

### *C. elegans* Strains and Maintenance

*C. elegans* were cultured using standard techniques at 20°C^23^. The following strains were used: wild type (WT) N2 Bristol, SPC321 [*alh*-*6(lax105)*], CL2166[*gst*-*4p::gfp*], SPC223 [*alh*-*6(lax105);gst*-*4p::gfp*], SPC542 [*alh*-*6(lax917);gst*-*4p::gfp*], SPC531 [*alh*-*6(lax906);gst*-*4p::gfp*], SPC528 [*alh*-*6(lax903);gst*-*4p::gfp*], SPC552 [*alh*-*6(lax927);gst*-*4p::gfp]*, SPC561 [*alh*-*6(lax937);gst*-*4p::gfp*], SPC564 [*alh*-*6(lax940);gst*-*4p::gfp*], SPC549 [*alh-6(lax924);gst*-*4p::gfp*], SPC566 [*alh*-*6(lax945);gst*-*4p::gfp*], SPC540 [*alh*-*6(lax915);gst*-*4p::gfp*], SPC562 [*alh*-*6(lax938);gst*-*4p::gfp]*, SPC563 [*alh*-*6(lax939);gst*-*4p::gfp*], SPC546 [*alh*-*6(lax921);gst*-*4p::gfp*], SPC527 [*alh*-*6(lax902);gst*-*4p::gfp*], SPC529 [*alh*-*6(lax904);gst*-*4p::gfp*], SPC536 [*alh*-*6(lax911);gst*-*4p::gfp*], SPC534 [*alh*-*6(lax909);gst*-*4p::gfp*], SPC559 [*alh*-*6(lax935);gst*-*4p::gfp*], SPC532 [*alh*-*6(lax907);gst*-*4p::gfp*], SPC569 [*alh*-*6(lax993);gst*-*4p::gfp*], SPC544 [*alh*-*6(lax919);gst*-*4p::gfp*], SPC562 [*alh*-*6(lax938);gst*-*4p::gfp*], SPC551 [*alh*-*6(lax926);gst*-*4p::gfp*], SPC530 [*alh*-*6(lax905);gst*-*4p::gfp*], SPC533 [*alh*-*6(lax908);gst*-*4p::gfp*], SPC548 [*alh*-*6(lax923);gst*-*4p::gfp*], SPC550 [*alh*-*6(lax925);gst*-*4p::gfp*], SPC565 [*alh*-*6(lax941);gst*-*4p::gfp*], SPC538 [*alh*-*6(lax913);gst*-*4p::gfp*], SPC543 [*alh*-*6(lax918);gst*-*4p::gfp*], SPC541 [*alh*-*6(lax916);gst*-*4p::gfp*], SPC545 [*alh*-*6(lax920);gst*-*4p::gfp*], SPC554 [*alh-6(lax929);gst*-*4p::gfp*], SPC558 [*alh*-*6(lax934);gst*-*4p::gfp*], SPC526 [*alh*-*6(lax901);gst*-*4p::gfp*], SPC568 [*alh*-*6(lax992);gst*-*4p::gfp*], SPC535 [*alh*-*6(lax910);gst*-*4p::gfp*], SPC553 [*alh*-*6(lax928);gst*-*4p::gfp*], SPC539 [*alh*-*6(lax914);gst*-*4p::gfp*], SPC556 [*alh*-*6(lax932);gst*-*4p::gfp*], SPC557 [*alh*-*6(lax933);gst*-*4p::gfp*], SPC567 [*alh*-*6(lax947);gst*-*4p::gfp*].

Double mutants were generated by standard genetic techniques. *E. coli* strains used were as follows: *OP50/E.coli* B for standard growth. All genetic mutants were backcrossed at least 4X prior to phenotypic analyses.

### Genetic Complementation (cis-trans) Testing

Hermaphrodites from each isolated mutant that phenocopied the *alh*-*6(lax105)*-like, age-related activation of the *gst*-*4p::gfp* reporter in the musculature were mated to SPC223 [*alh*-*6(lax105);gst*-*4p::gfp*] males. F1 progeny were screened at day 3 of adulthood for the *alh*-*6(lax105)*-like phenotype, which indicates a failure of the *alh*-*6(lax105)* allele to complement the mutation in the new mutant strain; thus the new mutant harbors a loss-of-function allele in *alh*-*6*.

### DNA Sequencing of *alh*-*6* Genetic Mutants

Approximately 200 adult worms were collected and washed with M9. Animals were homogenized and genomic DNA was extracted using the Zymo Research Quick-DNA Miniprep kit (Cat. #D3025). The entire *alh*-*6* genomic sequence (ATG to stop) was amplified by PCR and cloned in a linearized pMiniT 2.0 vector (NEB PCR Cloning Kit, Cat. #E1202S). Plasmid DNA was purified using the Zymo Research Zyppy Plasmid Miniprep kit (Cat. D4019) and sequenced.

### Microscopy

Zeiss Axio Imager and ZEN software were used to acquire all images used in this study. For GFP reporter strains, worms were mounted in M9 with 10mM levamisole and imaged with DIC and GFP filters. Worm areas were measured in ImageJ software (National Institutes of Health, Bethesda, MD) using the polygon tool.

### HRS Human Samples

The Health and Retirement Study (HRS^2,3^) is a nationally representative, longitudinal sample of adults aged 50 years and older, who have been interviewed every two years, beginning in 1992. Because the HRS is nationally representative, including households across the country and the surveyed sample now includes over 36,000 participants, it is often used to calculate national prevalence rates for specific conditions, including physical and mental health outcomes, cognitive outcomes, as well as financial and social indicators.

The sample for the current study is comprised of a subset of the HRS for which genetic data were collected, as described below. To reduce potential issues with population stratification, the GeneWAS in this study was limited to individuals of primarily European ancestry. The final subsample varied depending on the phenotype assessment. The subsample for each phenotype scan ranged from N=3,319 (for change in gait speed) to N=9,886 (for arm lifting), with the proportion of women at 58.5%.

### Genotyping Data

For HRS, genotype data were accessed from the National Center for Biotechnology Information Genotypes and Phenotypes Database (dbGaP^4^). DNA samples from HRS participants were collected in two waves. In 2006, the first wave was collected from buccal swabs using the Qiagen Autopure method (Qiagen, Valencia, CA). In 2008, the second wave was collected using Oragene saliva kits and extraction method. Both waves were genotyped by the NIH Center for Inherited Disease Research (CIDR; Johns Hopkins University) using the HumanOmni2.5 arrays from Illumina (San Diego, CA). Raw data from both phases were clustered and called together. HRS followed standard quality control recommendations to exclude samples and markers that obtained questionable data, including CIDR technical filters^24^, removing SNPs that were duplicates, had missing call rates ≥ 2%, > 4 discordant calls, > 1 Mendelian error, deviations from Hardy-Weinberg equilibrium (at *p*-value < 10^−4^ in European samples), and sex differences in allelic frequency ≥ 0.2). Further detail is provided in HRS documentation^18^. Applying these criteria to the gene region, on chromosome 1, (NC_000001.10): 19,194,787 - 19,232,430 resulted in available data on 53 SNPs. Because the goal is to evaluate whether SNPs within the same gene are associated with the phenotypes of interest, SNPs were not filtered for degree of linkage disequilibrium.

### Statistical Analysis of HRS Data Set

Following SNP extraction, we followed two analytical steps: (1) Phenotype construction, (2) Gene-wide association scans (GeneWASs), and (3) SNP comparisons.

### HRS Phenotype Construction

HRS phenotype construction was completed to calculate age-related ability or decline in ability over time. Table 2 shows the HRS data years from which phenotypes were calculated and details on how the variable is defined, and score or variable range. Datasets from multiple survey years were merged to get repeated assessments of variables on the same individuals. Phenotypes were calculated based on consensus following a review of the literature on assessments for age-related outcomes for variables implemented in the HRS and similar population-based surveys of aging. Further background for coding of specific phenotypes are described in detail previously for gait speed^25^ and for grip strength^25–27^. Phenotypes reflecting age-related ability were calculated at single timepoints, taking the most recent assessment per individual, assessed in HRS between the years of 2006-2012. Seven phenotypes were assessed by determining whether individuals had difficulty performing the activity. These include arm lifting, getting up from a chair, lifting 10 pounds, walking across the room, walking one block, walking several blocks, or jogging one mile. Two phenotypes, grip strength and gait speed, were assessed as performance on a measured task. Two phenotypes, grip strength decline and gait speed decline, were assessed as change in performance on those tasks over time. Five phenotypes, represented by composite scores listed in Supplementary Table 2, reflect age-related decline in ability and were calculated as change scores. All phenotypes reflecting change were calculated by taking the score from the most recent assessment and subtracting the score from the first assessment for each person, within the respective years listed. Additional descriptive statistics on phenotypes can be provided. Phenotypes were calculated using SAS 9.4.

### GeneWAS

GeneWAS occurred through a series of separate linear or logistic regression scans, under an additive model, adjusting for relevant covariates and indicators of population stratification as described below.

#### Population Stratification

As with any statistical analysis of association, if the correlation between dependent and independent variables differs for subpopulations, this may result in spurious genetic associations^28^. To reduce such type 1 error, we conducted the GeneWAS adjusting for population structure as indicated by latent factors from principal components analysis (PCA)^29,30^. Detailed descriptions of the processes employed for running PCA, including SNP selection, are provided by HRS^15^, and follow methods outlined by Patterson and colleagues)^30^. Two PCAs were run. The first PCA included 1,230 HapMap anchors from various ancestries and were used to test against self-reported race and ethnic classifications. Several corrections to the dataset were made based on this analysis. The second PCA was run on the corrected dataset, on unrelated individuals and excluding HapMap anchors, to create eigenvectors to serve as covariates and adjust for population stratification in association tests. From the second PCA, the first two eigenvalues with the highest values accounted for less than 4.5% of the overall genetic variance, with additional components (3-8) increasing this minimally, by a total of ~1.0%^18^. Based on these analyses, we opted for a strategy that does not ignore population substructure, but also does not over-correct, and adjusted for the first four PCs in all analyses. When coupling this approach of adjusting for principal components with all quality control procedures performed, excluding any related individuals and limiting the dataset for ancestral homogeneity, we reduce the likelihood of false associations resulting from population stratification)^29–35^.

#### Regression models and other covariates

When conducting regressions on phenotypes indicating change over time, additional adjustments were made using covariates for baseline levels, number of years during which change was calculated, and variables shown to affect outcomes. For example, with change in gait speed, a linear regression scan was run adjusting for sex, age at the first assessment point, number of years of followup, baseline walking speed, and floor type in addition to principal components. All GeneWAS were completed using PLINK 2.0^36^. The strength of the associations, as indicated by effect sizes and *p*-values, are not directly comparable for each phenotype because the sample sizes differed by phenotype. Thus, the strength of an association does not reflect how strong a SNP effects one phenotype compared to another. Also, because samples sizes were smaller for some phenotypes, we were less likely to detect true associations that may exist. However, because we did find multiple variants associated with certain phenotypes that were available for smaller sub-samples (e.g., N=3319 for gait speed decline), we are more confident that these results were not due to type 1 error. Because some phenotypes were measured with a dichotomous outcome (yes vs. no for having difficulty), we could not assess decline in ability over age for these phenotypes.

### SNP Comparison

We evaluated SNP associations in the GeneWAS by p-value, using the genome-wide significance level corrected for multiple testing, at a *p*-value threshold of 0.003. Given that the GeneWAS was run based on hypotheses generated from experimental models on the association of gene variation with aging-related muscle functioning, we calculate the multiple test threshold as a Bonferroni correction for 16 phenotypes (0.05/16=0.003). We also discuss additional findings at the “suggestive association” level of *p*-value ≤ 0.013 such that results would be less subject to Type II error at the cost of an overly stringent Type I cutoff given the a priori hypotheses and smaller sample sizes available for some phenotypes. To compare SNP associations, we: 1) assessed the independent contribution of each SNP meeting genome-wide significance; and 2) compared effect sizes and *p*-values across the gene to identify patterns of overlap for SNPs associated with more than one phenotype. For visualization of the SNP plots, we used R (CRAN; https://www.r-project.org).

### WormLab Measurements

As previously described^22^, but in brief; 15-20 animals were moved to a NGM stock plate without *E. coli* OP50 and recorded in WormLab software (MBF Bioscience) for 2 minutes.

### Statistical Analysis of *alh*-*6* Genetic Mutants

Data are presented as mean ± SD. Comparisons and significance were analyzed in GraphPad Prism 8. Comparisons between more than two groups were done using ANOVA.

