## Supplemental Figures and Tables for "Genetic variation in *ALDH4A1* predicts muscle health over the lifespan and across species"

### SUPPLEMENTARY TABLES AND FIGURE LEGENDS

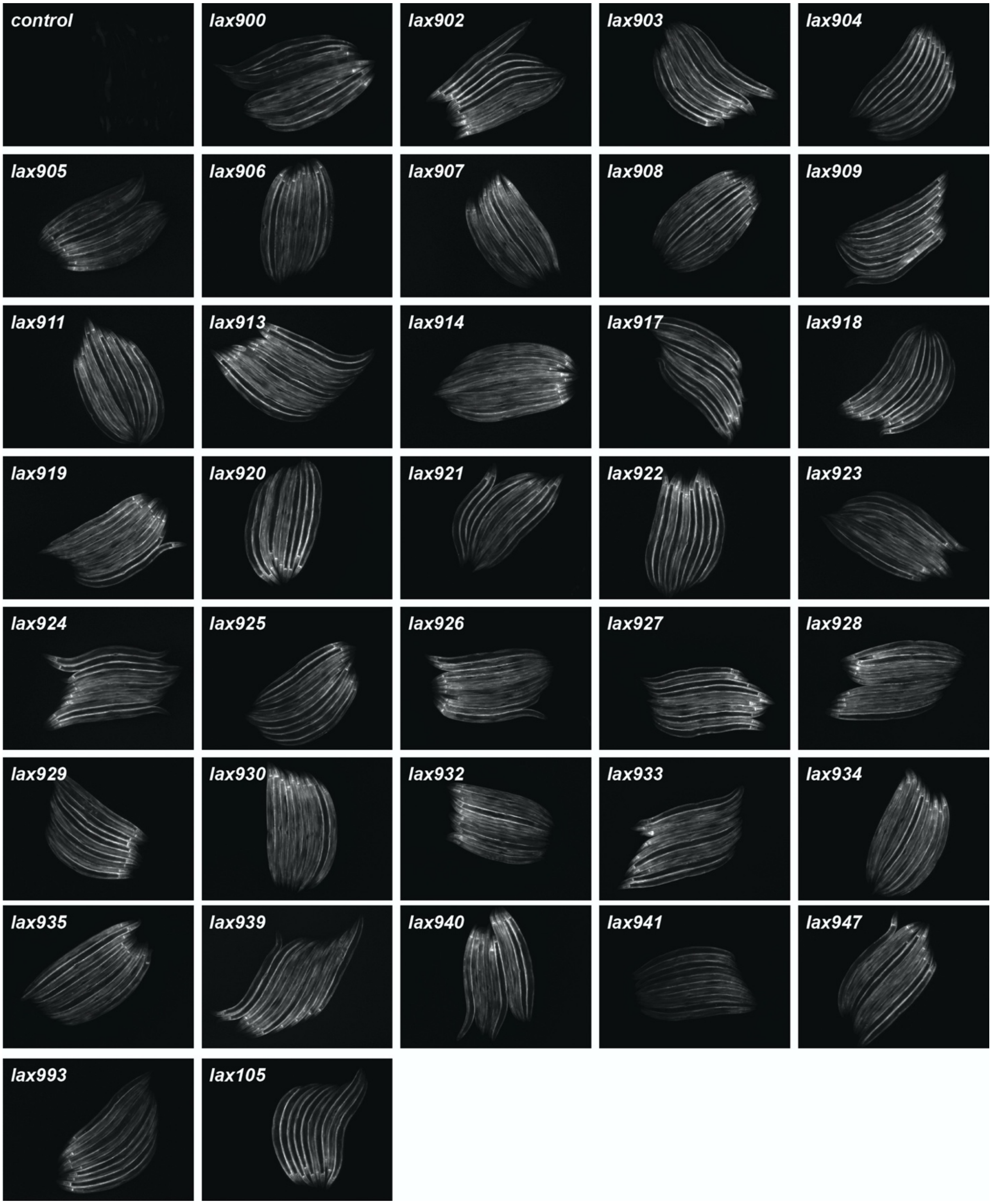


**Figure S1. Novel alleles of *alh-6* induce muscle specific activation of the SKN-1 reporter**. GFP fluorescence images of *gst-4p::gfp* animals harboring *alh-6* mutations, as indicated. 5X magnification. Quantified in Figure 1c.

**Table S1.** Details for phenotypes calculated from the U.S. Health and Retirement Study

| Variable | Ability or Decline | Scoring | Years Included | Assessment |
|---|---|---|---|---|
| **Individual phenotypes** | | | | |
| Grip strength decline over time | Decline | Change in kg of weight per year | 2006-2012 | Average performance when squeezing a hand dynamometer with the dominant hand twice |
| Grip strength ability | Ability | Kilograms of weight | 2006-2012 | Average performance when squeezing a hand dynamometer with the dominant hand twice |
| Arm Lifting | Ability | Difficulty (none vs some) | 2006- 2016 | Difficulty with reaching/extending arms above shoulder level |
| Getting up from a chair | Ability | Difficulty (none vs some) | 2006- 2016 | Difficulty with getting up from a chair after sitting for long periods |
| Lifting or carrying 10 lbs. | Ability | Difficulty (none vs some) | 2006- 2016 | Difficulty with lifting or carrying weights over 10 pounds |
| Gait Speed decline over time | Decline | Change in m/s per year | 2006-2012 | Average of walking performance over 2.5 meter walk |
| Gait Speed ability | Ability | Meters/second | 2006-2012 | Average of walking performance over 2.5 meter walk |
| Walking across a room | Ability | Difficulty (none vs some) | 2006- 2016 | Difficulty with walking across a room |
| Walking 1 block | Ability | Difficulty (none vs some) | 2006- 2016 | Difficulty with walking 1 block |
| Walking several blocks | Ability | Difficulty (none vs some) | 2006- 2016 | Difficulty with walking several blocks |
| Jogging a mile | Ability | Difficulty (none vs some) | 2006- 2016 | Difficulty with running or jogging about a mile |
| **Composite Indices** | | | | |
| Mobility Decline over time | Decline | Change in level of difficulty experienced over time | Between 1994-2016 | The Mobility score includes 5 tasks: walking several blocks, walking one block,walking across the room, climbing several flights of stairs and climbing one flight of stairs. |
| Large Muscle Group Decline | Decline | Change in level of difficulty experienced over time | Between 1994- 2016 | The Large Muscle Group score include 4 tasks: sitting for two hours, getting up from a chair, stooping or kneeling or crouching, and pushing or pulling a large object |
| Activities of Daily Living (ADL) Decline | Decline | Change in level of difficulty experienced over time | Between 1994- 2016 | ADL includes five tasks: bathing, eating, dressing, walking across a room, and getting in or out of bed |
| Instrumental Activities of Daily Living (IADL) Decline | Decline | Change in level of difficulty experienced over time | Between 1994- 2016 | IADL1 includes 3 tasks: using a telephone, taking medication, and handling money |

| | | | | |
|---|---|---|---|---|
| IADL2 Decline, expanded variable | Decline | Change in level of difficulty experienced over time | Between 1994- 2016 | IADL2 includes 5 tasks: using a telephone, taking medication, handling money, shopping, preparing meals |

**Table S2.** Top SNPs associated with multiple phenotypes at the significant and suggestive levels

| SNP | Phenotypes |
|---|---|
| rs111289603 (3'UTR) | Walking across a room, Walking 1 block, Getting up from a Chair, IADL decline, IADL2 decline (expanded measure) |
| rs28665699 (intron) | Grip strength decline, mobility decline |
| rs28493067 (intron) | Lifting or carrying 10 pounds, Mobility decline |
